# Induction of SARS-CoV-2 protein S-specific CD8+ T cells in the lungs of gp96-Ig-S vaccinated mice

**DOI:** 10.1101/2020.08.24.265090

**Authors:** Eva Fisher, Laura Padula, Kristin Podack, Katelyn O’Neill, Matthew M Seavey, Padmini Jayaraman, Rahul Jasuja, Natasa Strbo

**Affiliations:** Department of Microbiology and Immunology, Miller School of Medicine, University of Miami, Miami, FL, USA; Heat Biologics, Inc. Morrisville, NC, USA

**Keywords:** heat shock protein, gp96, vaccine, lungs, COVID-19, SARS-CoV-2 protein S, CD8+ T cells

## Abstract

Given the aggressive spread of COVID-19-related deaths, there is an urgent public health need to support the development of vaccine candidates to rapidly improve the available control measures against SARS-CoV-2. To meet this need, we are leveraging our existing vaccine platform to target SARS-CoV-2. Here, we generated cellular heat shock chaperone protein, glycoprotein 96 (gp96), to deliver SARS-CoV-2 protein S (spike) to the immune system and to induce cell-mediated immune responses. We showed that our vaccine platform effectively stimulates a robust cellular immune response against protein S. Moreover, we confirmed that gp96-Ig, secreted from allogeneic cells expressing full-length protein S, generates powerful, protein S polyepitope-specific CD4+ and CD8+ T cell responses in both lung interstitium and airways. These findings were further strengthened by the observation that protein-S -specific CD8+ T cells were induced in human leukocyte antigen (HLA)-A2-02-01 transgenic mice thus providing encouraging translational data that the vaccine is likely to work in humans, in the context of SARS-CoV-2 antigen presentation.

## Introduction

The rapid spread of the global COVID-19 pandemic has put pressure on the development of a SARS-CoV-2 vaccine to address global health concerns. We generated a gp96-Ig-secreting vaccine expressing full-length spike or “S” glycoprotein of SARS-CoV-2 via a cell-delivered platform. Targeting SARS-CoV-2 spike (S) protein remains the favorable vaccine choice as it is one of the most abundant and immunogenic proteins translated from the SARS-CoV-2 genome.^1^ Antibodies targeting S protein aim to neutralize mammalian host-cell interaction, thereby minimizing viral multiplicity of inflection, however, recent studies have shown that “antibodies are not enough” to protect against COVID-19 for a variety of reasons, including S-protein glycosylation, which shields the antibody from eliciting an optimal neutralization response.^2^ Antibody decay has also been detected in individuals after recovery from COVID-19, and this decline was more rapid than reported for the first SARS infection in 2003.^3,4^

T-cell immunity plays a pivotal role in generating a durable, immune memory response to protect against viral infection. Prior studies have shown that memory B-cell responses tend to be short lived after infection with SARS-CoV-1.^5,6^ In contrast, memory T-cell responses can persist for many years.^7^ Recent data confirm that SARS-CoV-2-specific memory CD8+ T cells are present in the vast majority of patients following recovery from COVID-19,^7-10^ and their protective role has been inferred from studies in patients who have had both SARS and MERS.^11-13^ Recent reports show that patients who have recovered from a severe SARS-CoV-2 infection have T-cell responses against viral spike protein and other structural and nonstructural proteins; in some patients, T-cell responses were present regardless of symptoms or antibody seropositivity.^14-16^ Here, we generated a COVID-19 vaccine based on the proprietary secreted heat shock protein, gp96-Ig vaccine strategy, that induces antigen-specific CD8+ T lymphocytes in epithelial tissues, including lungs.

Tissue-resident memory T (TRM) cells have been recognized as a distinct population of memory cells that are capable of rapidly responding to infection in the tissue, without requiring priming in the lymph nodes.^17-20^ Several key molecules important for CD8+ T cell entry and retention in the lung have been identified^21-26^ and recently CD69 and CXCR6^20,27-29^ have been confirmed as core markers that define TRM cells in the lungs. Furthermore, it was confirmed that CXCR6-CXCL16 interactions control the localization and maintenance of virus-specific CD8+ TRM cells in the lungs.^20^ It has also been shown that, in heterosubtypic influenza challenge studies,^30-32^ TRM were required for effective clearance of the virus. Therefore, vaccination strategies targeting generation of TRM and their persistence may provide enhanced immunity compared with vaccines that rely on circulating responses.^32^

Our platform technology consists of a genetically engineered construct of gp96, fusion protein gp96-Ig, wherein the C-terminal KDEL-retention sequence was replaced with the fragment crystallizable (Fc) portion of immunoglobulin G1 (IgG1), and then encoded within a plasmid vector that is transfected into a cell line of interest. The cell serves as the antigen supply to secreted gp96-Ig. Complexes of gp96-Ig and antigenic peptides lead to specific cross-presentation of cell-derived antigens by gp96-Ig in vivo.^33,34^ A crucial advantage offered by this gp96-based technology platform is that it allows for any antigen (such as SARS-CoV-2 S peptides) in the complex with gp96 to drive a potent and long-standing immune response. Over the last 2 decades, we have established that gp96-Ig, secreted from allogeneic or xenogeneic cells containing selected infectious antigens, generates potent, disease antigen specific, polyepitope, multifunctional CD8+ T cells in epithelial tissues.^33-39^ Here, we generated a COVID-19 vaccine based on the secreted heat shock protein, gp96-Ig vaccine strategy, and demonstrated vaccine-induced SARS-CoV-2 protein S-specific, CD8+ and CD4+ T lymphocytes in epithelial tissues, including lungs and airways. The secreted gp96-Ig-COVID-19 vaccine has the potential to elicit robust long-term memory T-cell responses against multiple SARS-CoV-2 antigens and is designed to work cohesively with other treatments/vaccines (as boosters or as second-line defense) with large-scale manufacturing potential.

## Methods

### Generation of Vaccine Cell Lines

Human embryonic kidney (HEK)-293 cells, obtained from the American Tissue Culture Collection (ATCC, #CRL-1573) and human lung adenocarcinoma cell lines (AD100)^40,41^ (source: University of Miami, FL, USA) were transfected with 2 plasmids: B45 encoding gp96-Ig (source: University of Miami) and pcDNA™ 3.1(-) (Invitrogen), encoding full-length SARS-CoV-2 protein S gene (Genomic Sequence: NC_045512.2; NCBI Reference Sequence: YP_009724390.1 GenBank Reference Sequence: QHD43416). The B45 plasmid expressing secreted gp96-Ig has been approved by the Food and Drug Administration and Office of Biotechnology Activities for human use and is currently employed in a clinical study for the treatment of nonsmall cell lung cancer (NSCLC) (NCT02117024, NCT02439450).^42^ The histidinol-selected, B45 plasmid, replicates as multicopy episomes and provides high levels of expression. Full-length SARS-CoV-2 protein S is based upon published SARS-CoV-2 protein S sequence from the original Wuhan strain (GenBank Reference Sequence: QHD43416) and cloned into the neomycin-selectable eukaryotic expression vector, pcDNA 3.1(-). HEK-293 and AD100 cells were simultaneously transfected with B45 and pcDNA 3.1 plasmid by lipofectamine (Invitrogen) following the manufacturers’ protocols. Transfected cells were selected with 1 mg/mL of G418 (Life Technologies, Inc.) and with 7.5 mM of L-Histidinol (Sigma Chemical Co., St. Louis, MO, USA). After stable transfection, cell line was established, single cell cloning by limiting dilution assay was performed, and all the cell clones were first screened for gp96-Ig production and then for protein S expression. Vaccine cells sterility testing and IMPACT™ II polymerase chain reaction evaluation was performed for: Ectromelia, mouse rotavirus (EDIM), lymphocytic choriomeningitis virus (LCMV), lactate dehydrogenase-elevating virus (LDEV), mouse adenovirus (MAV1, MAV2), mouse cytomegalovirus (MCMV), mouse hepatitis virus (MHV), murine norovirus (MNV), mouse parvovirus (MPV), minute virus of mice (MVM), mycoplasma pulmonis, Mycoplasma sp., Polyoma, pneumonia virus of mice (PVM), Reovirus 3 (REO3), Sendai, Theiler’s murine encephalomyelitis virus (TMEV), and all test results were negative.

### Western Blotting and Enzyme-Linked Immunosorbent Assay (ELISA)

Protein expression was verified by SDS-page and Western blotting using rabbit anti-SARS-CoV-2 spike glycoprotein antibody (MBS 150780) at 1/1000 dilution and secondary antibody: Peroxidase AffiniPure F(ab’)_2_ Fragment Donkey Anti-Rabbit IgG (H+L) (Jackson ImmunoResearch Laboratories) horseradish peroxidase conjugated anti-rabbit IgG (Jackson ImmunoResearch) at 1/10,000 dilution. S protein was visualized by an enhanced chemiluminescence detection system (Amersham Biosciences, Piscataway, NJ, USA) (**Figure 1c**). Recombinant human coronavirus SARS-CoV-2 spike glycoprotein S1 (Fc Chimera) (ab272105, Abcam) was used as positive control (loaded 2.4 ug/lane). One million cells were plated in 1 mL for 24 hours and secreted gp96-Ig production was determined by ELISA using antihuman IgG antibody for detection and human IgG1 as a standard (**Figure 1b**).

**Figure 1:**
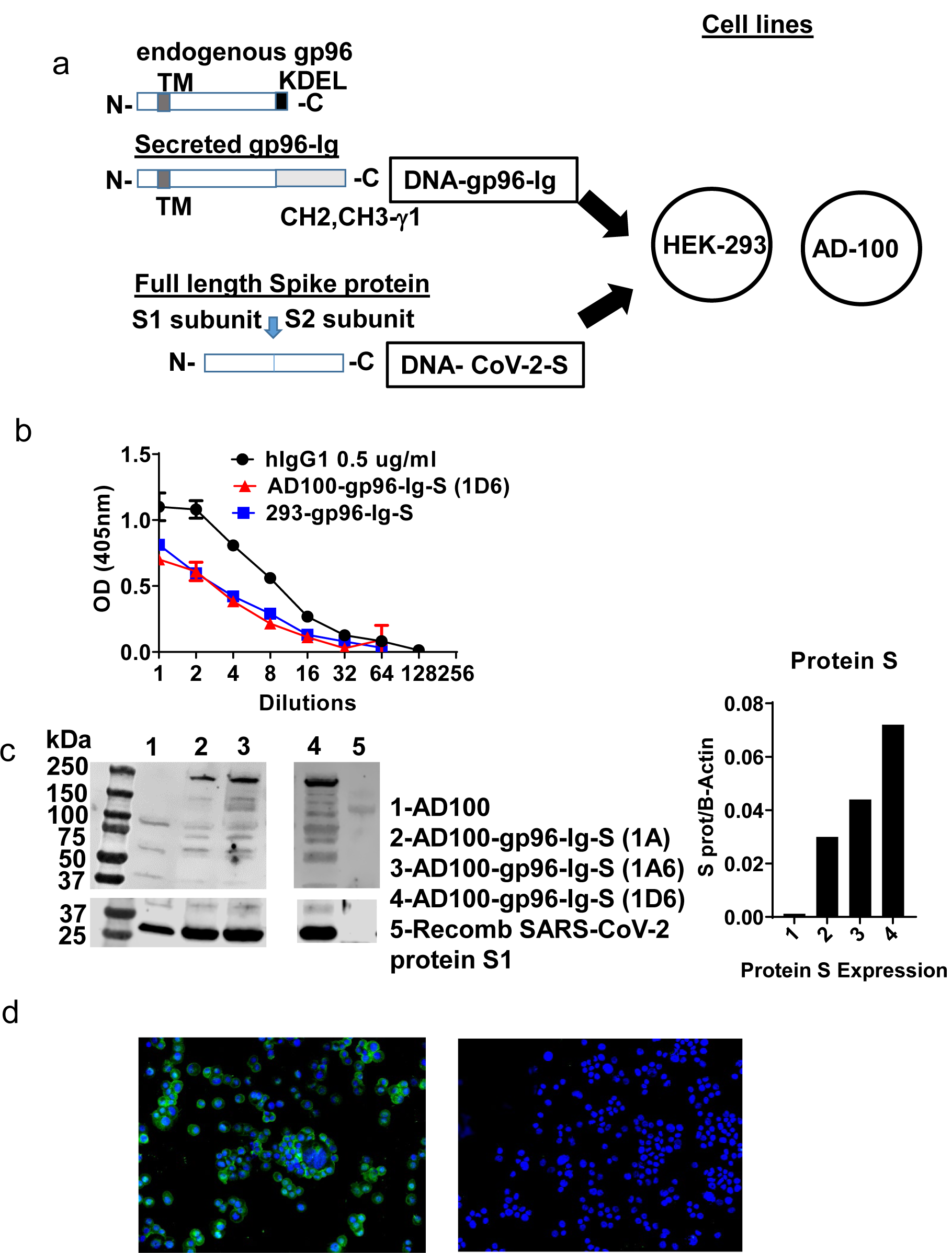
Schematic of gp96-Ig and SARS-CoV-2 protein S constructs used to generate vaccine cells HEK-293-gp96-Ig-S and AD-100-gp96-Ig-S. (a) Each panel presents the protein expressed by the DNA (black outline) for the gp96-Ig and SARS-CoV-2 protein S vaccine antigen. Gp96-Ig and SARS-CoV-2-S DNA were cloned into the mammalian expression vectors B45 and pcDNA 3.1, which are transfected into HEK-293 and AD100. Stable transfection vaccine cell clones (1A, 1A6, 1D6) were generated after selection with L-Histidinol and Neomycin; (b) One million 293-gp96-Ig-S and AD-100-gp96-Ig-S (1D6) cells were plated in 1 mL for 24 hours and gp96-Ig production in the supernatant was determined by ELISA using antihuman IgG antibody for detection with mouse IgG1 (0.5 ug/mL) as a standard; (c) Cell lysates were analyzed under reduced conditions by SDS-PAGE and Western blotting using anti protein S antibody and recombinant protein S1 as a positive control; (d) IF for protein S (in green) expressed in AD100-gp96-Ig-S cells using rabbit anti-SARS-CoV-2 S antibody and antirabbit Ig-AF488 as secondary antibody. AD100 was used as a negative control and β-actin for protein quantification. Original magnification 40× with DAPI nuclear staining shown in blue. DNA, deoxyribonucleic acid; ELISA, enzyme-linked immunosorbent assay; IgG, immunoglobulin G; N, amino terminus; C, carboxy terminus; IF, immunofluorescence; TM, transmembrane domain; KDEL, retention signal; CH2 CH3 gamma 1, heavy chain of IgG1. See text for explanation.

### Immunofluorescence (IF)

AD100-gp96-Ig cytospins were fixed in pure cold acetone (VWR chemicals, BDH^®^, Catalog #BDH1101) for 10 minutes followed by 3 washes of 5 minutes each with phosphate-buffered saline (PBS). The slides were left in blocking media (5% bovine serum albumin [BSA] in PBS) at room temperature for 2 hours. Rabbit anti-SARS-CoV-2 spike glycoprotein antibody (Abcam ab272504) and Donkey antirabbit IgG FITC, (BioLegend Cat# 406403) fluorescent antibody— were added in 1/50 and 1/100 dilutions of the antibodies combined in 5% BSA in PBS and/or rabbit isotype control (Abcam Ab172730 diluted 1/50), and incubated overnight at 4° C in a dark moisture chamber. The next day, slides were washed 3 times for 5 minutes with PBS and mounted with Prolong Gold antifade reagent with DAPI from Invitrogen (Catalog #36935), covered with a coverslip and allowed to cure. The slides were then sealed with nail polish and taken to the KEYENCE microscope for examination. The following filter cubes were used: DAPI (for nuclear stain), FITC (for protein S), and acquired on KEYENCE microscope (BZ-X Viewer).

### Animals and Vaccination

Mice used in this study were colony-bred mice (C57Bl/6) and human leukocyte antigen (HLA)-A02-01 transgenic mice (C57BL/6-Mcph1Tg (HLA-A2.1)1Enge/J, Stock No: 003475) purchased from JAX Mice (the Jackson Laboratory for Genomic Medicine, Farmington, CT, USA). Homozygous mice carrying the Tg (HLA-A2.1)1Enge transgene express human class I major histocompatibility complex (MHC) Ag HLA-A2.1. The animals were housed and handled in accordance with the standards of the Association for the Assessment and Accreditation of Laboratory Animal Care International under University of Miami Institutional Animal Care & Use Committee-approved protocol. Both female and male mice were used at 6–10 weeks of age.

Equivalent number of 293-gp96-Ig-protein S and AD100-gp96-Ig-protein S cells that produce 200-ng gp96-Ig or PBS were injected via the subcutaneous (s.c.) route in C57Bl/6 and HLA-A2 transgenic mice. Mice were sacrificed 5 days after vaccination and spleen, lungs, and bronchoalveolar lavage (BAL) were collected and processed into single-cell suspension.

### BAL and Lung Harvest and Cell Isolation

For mouse samples, spleens were collected, and tissues processed into single-cell suspension. Leukocytes were isolated from spleen and cervical lymph nodes by mechanical dissociation and red blood cells were lysed by lysing solution. BAL was harvested directly from euthanized mice via insertion of a 22-gauge catheter into an incision into the trachea. Hanks’ Balanced Salt Solution (HBSS) was injected into the trachea and aspirated 4 times. Recovered lavage fluid was collected and BAL cells were gathered after centrifugation.

To isolate intraparenchymal lung lymphoid cells, the lungs were flushed by 5 mL of prechilled HBSS into the right ventricle. When the color of the lungs changed to white, the lungs were excised avoiding the peritracheal lymph nodes. Lungs were then removed, washed in HBSS and cut into 300-mm pieces, and incubated in Iscove’s Modified Dulbecco’s Medium containing 1 mg/mL collagenase IV (Sigma) for 30 minutes at 37° C on a rotary agitator (approximately 60 rpm). Any remaining intact tissue was disrupted by passage through a 21-gauge needle. Tissue fragments and majority of the dead cells were removed by a 250-mm mesh screen, and cells were collected after centrifugation.

### Ex Vivo Stimulation and Intracellular Cytokine Staining

Spleen and intraparenchymal lung lymphocytes from immunized and control animals were analyzed for protein S-specific CD8+ T cell responses. 1-1.5×10/6 cells were incubated for 20 hours with 2 protein S peptide pools (S1 and S2, homologous to vaccine insert) (JPT Peptide Technologies, Berlin, Germany; PM-WCPV-S1). Peptide pools contain pools of 15-meric peptides overlapping by 11 amino acids covering the entire protein S proteins. Peptide pools were combined (S1+S2) and used at a final concentration of 1.25 ug/mL of each peptide, followed by addition of Brefeldin A (BD GolgiPlug™; BD Biosciences, San Diego, CA, USA) (10 ug/mL) for the last 5 hours of the incubation. Stimulation without peptides served as background control. The results were calculated as the total number of cytokine-positive cells with background subtracted. Peptide stimulated and non-stimulated cells were first labeled with live/dead detection kit (Thermo Fisher Scientific, Waltham, MA, USA) and then resuspended in BD Fc Block (clone 2.4G2) for 5 minutes at room temperature prior to staining with a surface-stain cocktail containing the following antibodies purchased from BioLegend^®^ (San Diego, CA, USA): antigen presenting cell (APC)Cy7 CD45; Clone; AF700 CD3: Clone: 17A2; APC CD4: Clone:RM4-5; PerCP CD8: Clone:53-6.7; PE Dazzle CD69: Clone:H1.2F3; BV 605 CD44: Clone:IM7; BV510 CD62L: Clone: MEL-14; PerCP/Cy5.5 CCR6 Clone: 29-2L17. After 30 minutes, cells were washed with a flow cytometry staining buffer and then fixed and permeabilized using BD Cytofix/Perm fixation/permeabilization solution kit (according to manufacturer instructions), followed by intracellular staining using a cocktail of the following antibodies purchased from BioLegend: Alexa Fluor 488 interferon (IFN) gamma: Clone: XMG1.2; PE interleukin 2 [IL-2]: Clone: JES6-5H4 PE Cy7 tumor necrosis factor alpha (TNFα): Clone: MPG-XT22.

Data were collected on Spectral analyzer SONY SP6800 instrument (Sony Biotechnologies, Inc, San Jose, CA, USA). Analysis was performed using FlowJo™ software version 10.8 (Tree Star Inc, Ashland, OR, USA). Cells were first gated on live cells and then lymphocytes were gated for CD3+ and progressive gating on CD8+ T cell subsets. Antigen-responding CD8 (cytotoxic) T cells (IFNγ, or IL-2, or TNFα-producing/expressing cells) were determined either on the total CD8+ T cell population or on CD8+ CD69+ cells.

### HLA-A02-01 Pentamer Staining

A total of 1-2×10^6^ spleen, BAL, or lung cells were labelled with peptide-MHC class I pentamer-APC (ProImmune, Oxford, UK) and incubated for 15 minutes at 37° C. Cells were labelled with LIVE/DEAD™ Fixable Violet – Dead Cell Stain Kit (Invitrogen, Carlsbad, CA, USA) and then stained with the following antibody cocktail: APCCy7 CD45; Clone; AF700 CD3: Clone: 17A2; PECy7 CD4: Clone:RM4-5; FITC and PerCP CD8: Clone:53-6.7; PE Dazzle CD69: Clone:H1.2F3; BV 605 CD44: Clone:IM7; BV510 CD62L: Clone: MEL-14; PerCP/Cy5.5 CCR6 Clone: 29-2L17; (clone). Spleen and lung cells that were stimulated overnight with peptide pools (as described under ex-vivo stimulation and intracellular staining) were fixed and permeabilized with Cytofix/Perm solution (BD) and then stained for intracellular cytokines: IFNγ, and IL-2. Cells were acquired on SP6800 Sony instrument and data analyzed using FlowJo software version 10.8. Data were analyzed using forward side-scatter single-cell gate followed by CD45, CD3, and CD8 gating, then pentamer gating within CD8+ T cells. These cells were then analyzed for expression of markers using unstained and overall CD8+ population to determine the placement of the gate. Single-color samples were run for compensation and fluorescence minus 1 control sample were also applied to determine positive and negative populations, as well as channel spillover.

### Statistics

All experiments were conducted independently at least 3 times on different days. Comparisons of flow cytometry cell frequencies were measured by the 2-way analysis of variance (ANOVA) test with Holm-Sidak multiple-comparison test, *p<0.05, **p<0.01, and ***p<0.001, or unpaired T-tests (2-tailed) were carried out to compare the control group with each of the experimental groups (alpha level of 0.05) using the Prism software (GraphPad Software, San Diego, CA, USA). Welch’s correction was applied with the unpaired T test, when the p-value of the F test to compare variances were ≤0.05. Data approximately conformed to Shapiro-Wilk test and Kolmogorov-Smirnov tests for normality at 0.05 alpha level. Data were presented as mean ± standard deviation in the text and in the figures. All statistical analysis was conducted using GraphPad Prism 8 software.

## Results

### AD100 and HEK-293 Express gp96-Ig and Protein S

Cell-based secreted heat shock protein technology has been previously validated in numerous animal models and in humans.^37-39,42^ The secreted form of gp96 protein (gp96-Ig) was generated by replacing the c-terminal, KDEL-retention sequence of human gp96 gene, with hinge region and constant heavy chains (CH2 and CH3) of human IgG1^43^ (**Figure 1a**). The pcDNA 3.1(–) vector was used to express SARS-CoV-2 spike (S) protein (in this manuscript referred as protein S) (**Figure 1a**), due to its propensity to constitutively express large amounts of the proteins in a mammalian cells. Complementary (c) DNA encoding the full-length SARS-CoV-S glycoprotein included Kozak sequence (GCCACC) to optimize expression in eukaryotic cells and the open-reading frame contained endogenous leader sequence, transmembrane, and cytosolic domains.

Vaccine cells, 293-gp96-Ig-S and AD100-gp96-Ig-S, were generated by cotransfection of AD100 and HEK293 cells with plasmids encoding gp96-Ig (B45) and protein S (pcDNA 3.1) and selection with G418 and L-histidinol as described in **Methods**. We confirmed by ELISA that both stable transfected cell lines secreted gp96-Ig into culture supernatants at a rate of 125 ng/mL/24 hours/10^6^ vaccine cells (**Figure 1b**). Our previous data indicate that gp96-Ig accumulation in cell culture supernatant is linear and time dependent.^35,43^

Protein S expression by the vaccine cells was confirmed by analyzing vaccine cell lysates on SDS page, blotting with anti-SARS-CoV-2 S antibody (**Figure 1c**) and by immunofluorescence (**Figure 1d**). We observed expression of full-length protein S (250 kDa) only in AD100 transfected cell lines (lanes 2–4) but not in nontransfected AD100 cell line (lane 1). In addition, we observed molecular weight bands of 120 and 130 kDa that could represent cleavage products of full length protein S (protein S1 and S2) and/or gp96-Ig fusion protein chaperoning the protein S peptides. The expected molecular weight of gp96-Ig fusion protein is 116 kDa.

Additional bands, of ∼70 kDa, were found to be expressed only in the transfected cell line and were not observed in the nontransfected AD100 cells. However, nonspecific bands of 100, 60, and 40 kDa were observed in the AD100 parental cell line. Recombinant protein S1 130 kDa was used as a positive control. We calculated the ratio of protein S to β-actin expression (**Figure 1c**) and confirmed the expression of protein S by immunofluorescence (**Figure 1d**). We observed cytoplasmic and transmembrane distribution of protein S in AD100-gp96-Ig-S cell line. We therefore confirmed the expression of gp96-Ig and S protein in our AD100 cell line and used it for immunogenicity studies as described below.

### Secreted gp96-Ig-S Vaccine Induces CD8+ T Cell Effector Memory (TEM) and TRM Responses in the Lungs

Our vaccination strategy is based on the quantity of gp96-Ig-S secreted by the vaccine cells to stimulate CD8+ CTL responses via APC cross-presentation. The vaccination dose, is therefore, standardized to a set amount of gp96-Ig secreted by 10^6^ vaccine cells within 24 hours. It has been well established from our previous vaccine immunogenicity studies that the optimal dose for induction of CD8+ T cell specific responses in mice is 200–500 ng/mL.^33,35,38,39,43^ Here, we used 200 ng/mL to immunize mice with AD100-gp96-Ig-S vaccine. Mice were vaccinated via the s.c. route and, after 5 days, the frequency of T cells within spleen, lungs (lung parenchyma), and BAL cells (lung airways) was determined. We observed significant increase in the frequencies of CD8+ T cells in the spleen and lungs, but not within the BAL of vaccinated mice (**Figure 2a**). Frequency of CD4+ T cells was unchanged between vaccinated and control mice in all analyzed tissues. It is well established that vaccination with gp96-Ig induces CD8+ TEM differentiation.^33,37,39^ Here, we confirmed that gp96-Ig-S vaccine primes strong effector memory CD8+ T-cell responses as determined by analysis of CD44 and CD62L expression (**Figure 2b**). Whereas the frequency of naïve (N), CD44-CD62L+CD8 T cells and central memory (CM), CD44+CD62L+ CD8+ T cells was unchanged, we found statistically significant increase of TEM CD44+CD62L-CD8+ T cells within the spleen and lungs (**Figure 2b**). In addition, we observed a trend of more TEM CD8+ T cells within the CD8+ T cells in the BAL (**Figure 2b**). TRM are a distinct memory T cell subset compared to CM and EM cells^44^ that are uniquely situated in different tissues, including lungs.^30,31^ One of the canonical markers of TRM T cells is CD69.^20,44,45^ We found that there was a significant increase in the frequency of CD8+CD69+ T cells in vaccinated mice compared to control, non-vaccinated mice in both spleen and lungs (**Figure 2c**). Even though the frequency of CD8+CD69+ T cells was the highest in the BAL compared to spleen and lungs, we did not observe a difference in their frequencies between vaccinated and control mice. Overall, vaccination with AD100-gp96-Ig-S induced robust TEM and TRM CD8+ T cell responses in both spleen and lungs. Our vaccine can therefore successfully elicit both systemic and tissue-specific immune response, which is pivotal in conferring robust immunity against infection such as against SARS-CoV-2.

**Figure 2:**
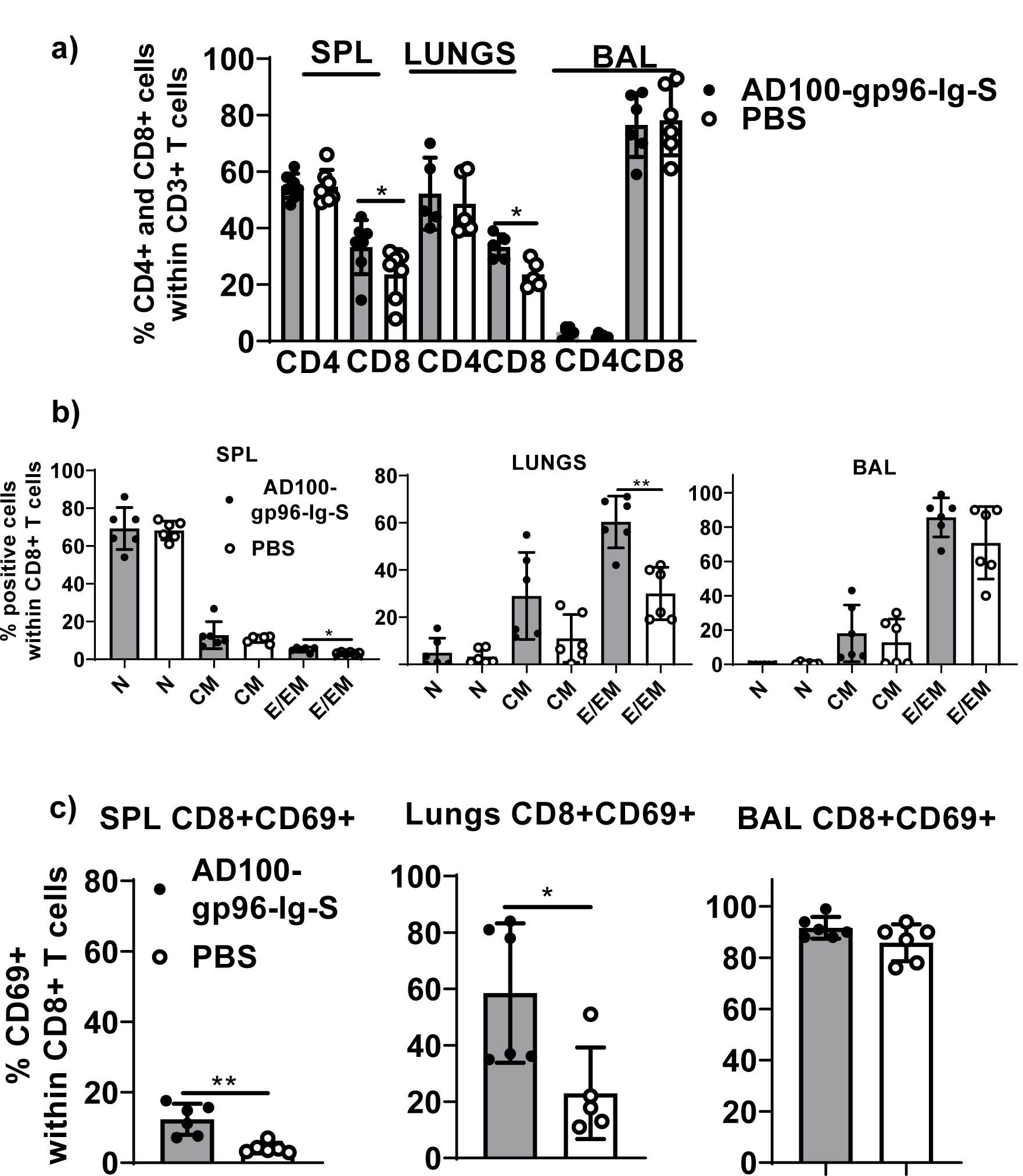
Secreted gp96-Ig-S vaccine induces CD8+ TEM and TRM responses in the lungs Equivalent number of AD100-gp96-Ig-S vaccine cells that produce 200 ng/mL gp96-Ig or PBS were injected by s.c. route in C56Bl/6 mice. 5 days later, mice were sacrificed and spleen, lungs, and BAL were isolated and (a) frequency of CD4+ and CD8+ T cells; (b) naive (N) CD44-CD62L+, CM CD44+CD62L+ and EM CD44+CD62L-CD8+ T cells and c) TRM CD69+ cells were determined by flow cytometry after staining the cells with antibodies against the following surface markers: CD45, CD3, CD4, CD8, CD44, CD62L and CD69 antibodies. Bar graph shows percentage of CD4+ and CD8+ cells within CD3+ cells or CD8+ T cell memory subset within CD8+ T cells. Data represent at least 2 technical replicates with 3-6 independent biological replicates per group. *p<0.05, **p<0.01, ***p<0.001. (a–b) Mann-Whitney tests were used to compare 2 experimental groups. To compare >2 experimental groups, Kruskal-Wallis ANOVA with Dunn’s multiple comparison tests were applied). BAL, bronchoalveolar lavage; CM, central memory; EM, effector memory; TEM, T cell effector memory; TRM, T cell resident memory.

### Both Protein S-Specific CD8+ and CD4+ T Helper 1 (Th1) T Cell Responses are Induced by gp96-Ig-S Vaccine

To evaluate polyepitope, protein S-specific CD8+ and CD4+ T-cell responses induced by gp96-Ig-S vaccination, we used pooled S peptides (S1+S2) and a multiparameter intracellular cytokine-staining assay to assess Th1 (IFNγ+, IL-2+ and TNFα+), CD8+ and CD4+ T cells (**Figure 3**). Spleen and lung cells were tested for responses to the pool of overlapping protein S peptides (S1 + S2) and all of the vaccinated animals showed significantly higher magnitude of the protein S-specific T cell responses against S1 and S2 epitopes compared with nonvaccinated controls (**Figures 3a–3d**). Increase in the vaccine-induced Th1 CD8 T cell responses (IFNγ+, IL-2+, and TNFα+) was noted in both spleen and lungs (**Figures 1a, 1b**), whereas Th1 CD4 T cell responses (IFNγ+, IL-2+, and TNFα+) were induced only in lungs (**Figures 3c, 3d**). The proportion of the protein S-specific CD8+ T cells that produce IFNγ (26.6%) was significantly reduced in the lungs (7%), while both TNFα and IL-2 productions were increased in the lungs (45% and 47%, respectively) compared to spleen (26% and 26%, respectively) (**Figure 3e**). We found that the proportion of the protein S-specific CD4+ T cells that produce IFNγ was higher in the spleen than in the lungs (57% [spleen] versus 27% [lungs]), whereas IL-2 production was higher in the lungs than in the spleen (15% [spleen] versus 34% [lungs]). (**Figure 3e**). Assessment of the polyfunctionality of protein S-specific CD8+ and CD4+ T cells in the spleen and lungs revealed that the vast majority of protein S-specific CD8+ and CD4+ T cells, irrespective of their location, synthesized only 1 cytokine (**Figure 3f**). Proportion of the protein S-specific CD8+ T cells in the spleen and lungs that produce 3 cytokines at the same time was higher than for CD4+ T cells. Only a small proportion of protein S-specific CD4+ T cells in the lungs produce 2 or 3 cytokines (3.6%, 2 cytokines; 1.5%, 3 cytokines) (**Figure 3f**).

**Figure 3:**
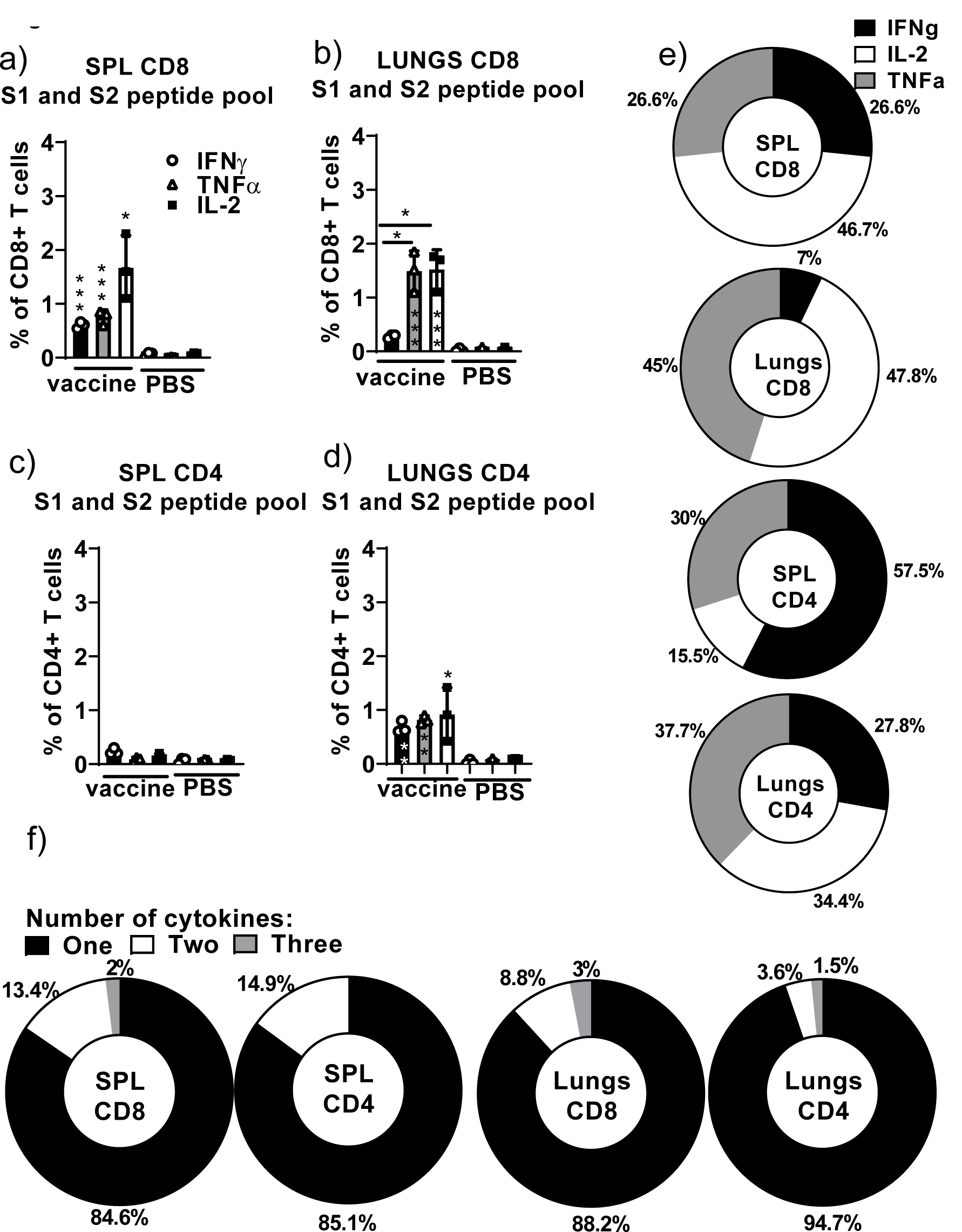
Secreted gp96-Ig-S vaccine induces protein S specific CD8+ and CD4+ T cells in the spleen and lung tissue. 5 days after the vaccination of C57Bl6 mice, splenocytes and lung cells were isolated from vaccinated and control mice (PBS) and in vitro restimulated with S1 and S2 overlapping peptides from SARS-CoV-2 protein in the presence of protein transport inhibitor, brefeldin A for the last 5 hours of culture. After 20 hours of culture, ICS was preformed to quantify protein S-specific CD8+ and CD4+ T-cell responses. Cytokine expression in the presence of no peptides was considered background and it was subtracted from the responses measured from peptide pool stimulated samples for each individual mouse. (a–b) CD8+ T cells from spleen and lungs expressing IFNγ, TNFα and IL-2 in response to S1 and S2 peptide pool; (c–d) CD4+ T cells from spleen and lungs expressing IFNγ, TNFα and IL-2 in response to S1 and S2 peptide pool; (e) Proportion of antigen (protein S)-experienced CD8+ and CD4+ T cells isolated from spleen and lung tissue expressing IFNγ, TNFα, or IL-2 after o/n stimulation with S1 + S2 peptides. Pie charts corresponding to cytokine profiles of CD8+ and CD4+ T cells isolated from spleen and lung tissue; (f) Polyfunctional profiles of antigen experienced CD8+ and CD4+ T cells. Pie charts corresponding to polyfunctional profiles of CD8+ CD4+ T cells isolated from spleen and lung tissue after o/n stimulation with S1 + S2 peptides. Assessment of the mean proportion of cells making any combination of 1–3 cytokines (IFN-γ, TNFα, IL-2). Data represent at least 2 technical replicates with 3–6 independent biologic replicates per group. *p<0.05, **p<0.01, ***p<0.001. Kruskal-Wallis ANOVA with Dunn’s multiple comparisons tests were applied. Asterisks (*) above or inside the column denote significant differences between indicated T cells producing cytokines in vaccine versus control (PBS) at 0.05 alpha level. ANOVA, analysis of variance; ICS, intracellular cytokine staining; IFN, interferon; IL, interleukin; PBS, phosphate-buffered saline; TNF, tumor necrosis factor.

It was therefore confirmed that a polyepitope, S-specific, polyfunctional CD4+ and CD8+ T cell response was generated in the spleen and lungs to different extents, providing a strong vaccine-induced Th1 cellular immune responses.

### Induction of SARS-CoV-2 Protein S Immunodominant Epitope-Specific CD8+ T Cells in the Lungs and Airways of Vaccinated HLA-A2-Transgenic Mice

Recently, it was reported that polyfunctional SARS-CoV-2-specific memory CD8+ T cell responses generated against cognate antigens positively correlated with a number of symptom-free days after infection.^14,16^ Therefore, it is important to develop vaccines that can elicit SARS-CoV-2-specific CD8+ T cells. Having identified overall T-cell responses to SARS-CoV-2 protein S (**Figure 3**), we wanted to determine whether gp96-Ig-S vaccine induced HLA class I-specific cross-presentation of immunodominant SARS-CoV-2 protein S epitopes. In order to do this, we used transgenic HLA-A 02:01 mice and HLA class I pentamers as probes to detect CD8+ T cells specific for 2 immunodominant SARS-CoV-2 protein S epitopes: YLQPRTFLL (YLQ) (aa 269-277) and FIAGLIAIV (FIA) (aa 1220-1228) in vaccinated mice (**Figure 4**). We found that the vaccine effectively induces both YLQ+CD8+ T cells, as well as FIA+CD8+ T cells in the spleen, lungs, and BAL (**Figure 4**). Interestingly, we found the highest frequency of YLQ+CD8+ T cells in the BAL of vaccinated mice and the lowest frequency of YLQ+ and FIA+ CD8+ T cells was observed in the lungs. Upon further phenotype analysis of YLQ+CD8+ T cells, it was confirmed that they express both CD69 and CXCR6 (**Figure 5**). Particularly, we found that all YLQ+CD8+ T cells in the BAL were also CXCR6+, and the frequency of YLQ+CD8+CXCR6+ cells was significantly higher in the BAL compared to the lungs.

**Figure 4:**
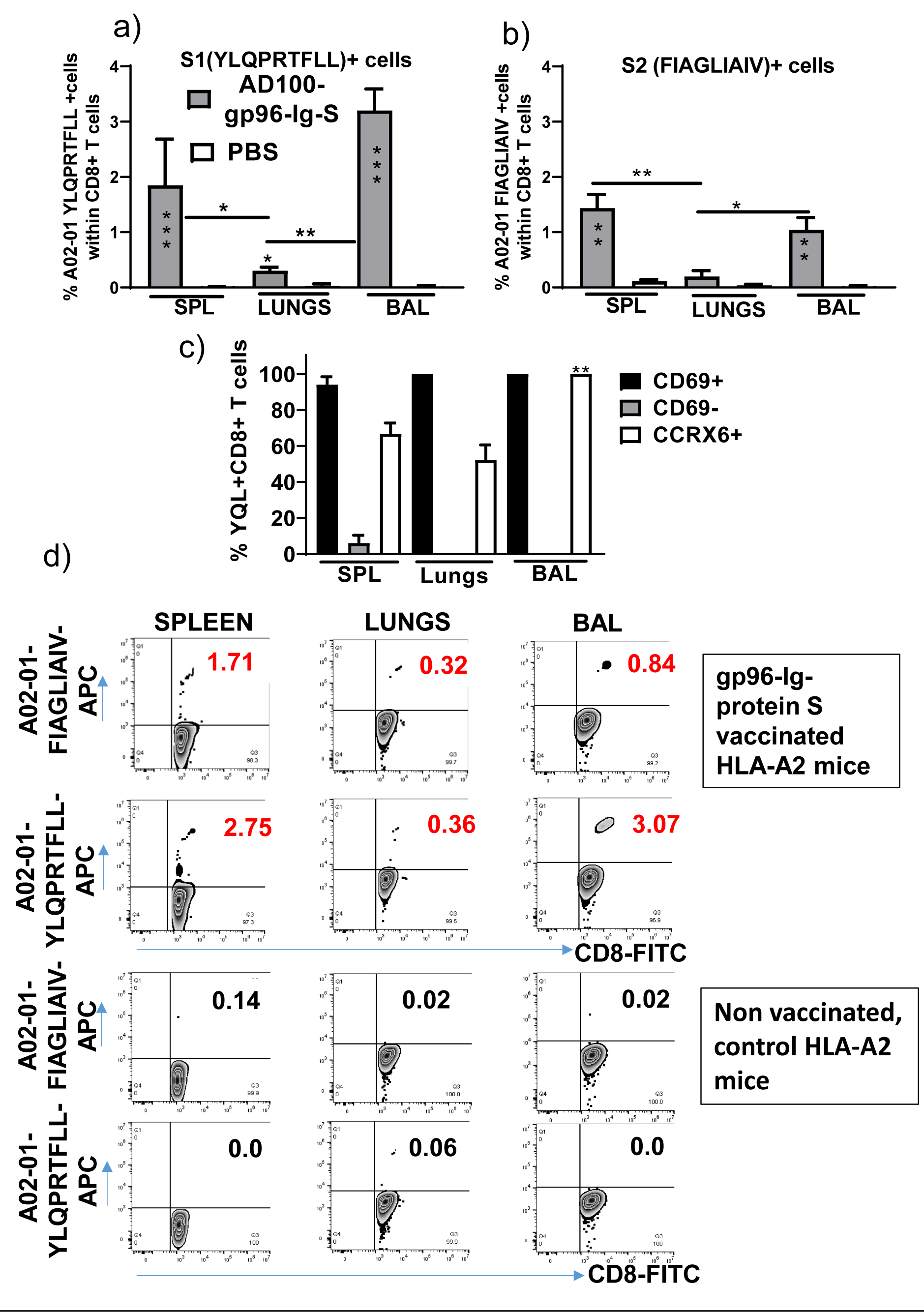
Secreted Gp96-Ig-S vaccine induces S1- and S2-specific CD8 + CD69 + CXCR6 + cells in the spleen, lung tissue, and BAL. 5 days after the vaccination of HLA-A2 transgenic mice, splenocytes, and lung cells were isolated form vaccinated and control mice (PBS). Cells were stained with HLA-A2 02-01 pentamer containing FIAGLIAIV and YLQPRTFLL peptides, followed by surface staining for CD45, CD3, CD4, CD8, CD69, CXCR6. (a–b) Bar graphs represent percentage of the pentamer positive cells within CD8+ T cells; (c) Bar graphs represent percentage of CD69+, CD69-, and CXCR6+ cells within YQL-pentamer positive cells; (d) Representative zebra plots of gated CD8+ T cells expressing indicated pentamer-specific TCR+ CD8+ T cells in vaccinated and nonvaccinated HLA-A2 mice. Data represent at least 2 technical replicates with 3–6 independent biologic replicates per group. *p<0.05, **p<0.01, ***p<0.001. Kruskal-Wallis ANOVA with Dunn’s multiple comparisons tests were applied. Asterisks (*) above or inside the column denote significant differences between indicated pentamer+CD8+ T cells in the vaccinated group and control (PBS) (a–b) and between BAL and lungs (c) at 0.05 alpha level. ANOVA, analysis of variance; BAL, bronchoalveolar lavage; PBS, phosphate-buffered saline.

## Discussion

Our vaccine approach is based on the gp96-Ig platform technology that elicits potent, antigen-specific CD8+ T-cells. This proprietary secreted heat shock protein platform has been successfully used to induce immunogenicity against tumors, HIV/SIV, Zika, and malaria in different animal models.^37-39,46-48^ Importantly, this vaccine strategy has shown success in delaying virus acquisition, as well as in improving the survival of NSCLC patients in clinical trials.^38,42^

The principle of a cell-based vaccine relies on the ability of gp96-Ig to chaperone antigenic proteins to be efficiently endocytosed and cross-presented by activated dendritic cells (DC) to CD8+ T cells, thereby stimulating an avid, pathogen-specific T-cell response.^33,34,37-39^ We adapted this cell-based technology to create a vaccine that delivers SARS-CoV-2 spike (S) protein directly to DCs, so that primed and activated SARS-CoV-2 protein S-specific CD8+ T cells can identify and kill SARS-CoV-2 infected lung epithelial cells. We generated vaccine cells by co-expressing secreted gp96-Ig and full-length protein S. Gp96-Ig is an endoplasmic reticulum chaperone that, together with TAP (transporter associated with antigen processing) and calreticulin in the endoplasmic reticulum, is thought to constitute a relay line for antigenic peptide transfer from the cytosol to MHC class I molecules in a concerted and regulated manner.^49,50^ The gp96-antigenic peptide complexes are predominantly internalized by subsets of APCs through cell surface receptor CD91. Internalized gp96 can effectively present the associated peptides to MHC class I and II molecules and thus activate specific CD8+ and CD4+ T-cell responses.^34,39,51,52^ We expressed full-length protein S in the vaccine cells to ensure broad representation of all immunodominant protein S peptides (S1- and S2-derived peptides) by secreted gp96-Ig. Since coronaviruses assemble in the compartment between the endoplasmic reticulum and Golgi apparatus^53,54^ and the S leader directs it to the endoplasmic reticulum, the native leader sequence of protein S was retained, as well as transmembrane and cytosolic domain. We confirmed in previous studies that secreted gp96-Ig provides immunologic specificity for the antigenic repertoire expressed inside of the cells, including surrogate antigen ovalbumin, as well as numerous tumor or infectious antigens, but does not cross-immunize to different cell-derived antigens.^35,37-39^ Our data are consistent with the explanation that S1 and S2 peptides associated with secreted gp96-Ig are transferred to and presented by class I and II MHC and stimulate a S1- and S2-specific CD8+ and CD4+ T cell response. We confirmed that vaccination with AD100-gp96-Ig-S induces CD8+ T cells specific for S1- and S2-immunodominant epitopes in both lungs and airways. Most importantly, this is a proof-of-concept study that will be applied to other structural proteins such as nucleocapsid protein, membrane protein, and nonstructural proteins such as NSP-7, NSP-13 of ORF-1 that all have been reported to be important in induction of SARS-CoV-2-specific CD4 and CD8 T cell responses in convalescents.^7,10,11,55^

In agreement with our previous findings,^33,38,39^ the gp96-Ig vaccine resulted in the preferential induction of CD8+ T cell responses systemically and in epithelial compartments. However, this is the first report about the increase in the frequencies of vaccine induced CD8+ T cells in the lungs (**Figure 1a**). TEM CD8+ T cells are considered to constitute the frontline defense within the different epithelial compartments including lungs and airways, which promptly recognize and kill infected cells. Our data suggest that after a single dose of AD100-gp96-Ig-S immunization there is preferential compartmentalization of TEM and TCM immune responses in the lungs and BAL compared to the spleen where a majority of cells are naïve CD8+ T cells (Fig 1b). However, additional memory cells without migratory potential such as TRM CD8+ T cells, exist within the tissues including lungs and airways.^20,44,45,56^ Since TRM are uniquely situated in the lungs to immediately respond to reinfection, by inducing the protein-S-specific CD8+ T cells that home to the lungs, gp96-Ig-S vaccine provides an ideally balanced generation of both arms, TRM and TEM, of the memory response in the lungs. To further gauge the effect of gp96-Ig vaccination on the induction of epitope specific immunogenicity, we used pentamers to detect S1 and S2 epitope-specific CD8+ T cell responses. We found that gp96-Ig induced the highest frequencies of S1- and S2-epitope specific CD8+ T cells in the airways. In light of the new findings about exclusive highly clonally expanded SARS-CoV-2-specific CD8+ T cells with preferentially expressed tissue-resident genes (XCL1, CXCR6 and ITGAE) in the BAL of moderate COVID-19 cases^57^ and not in the critical/severe COVID-19 patients, induction of SARS-CoV-2 specific CD8+ T cells that home to airway epithelium emphasizes the importance of developing vaccination strategies that induce TRM antigen-specific CD8+ T cells that will improve efficacy of vaccination against respiratory pathogens including SARS-CoV-2.

It is well appreciated that the antigen presenting cells at the site of immunization direct the imprinting of the ensuing T-cell response and control the expression of trafficking molecules.^58^ Priming of CD8+ T cells by CD103+ DC was found to promote TRM CD8+ T cell differentiation and migration into peripheral epithelial tissues, including lungs.^56,59^ Our previous studies indicated that gp96-Ig immunization increases frequency of CD11c^high^ MHC class II^high^ CD103+ cells at the vaccination site.^33^ In light of our previous findings and findings of Bedoui et al^60^ that CD103+ DCs are the main migratory subtype with dominant cross-presenting ability, induction of CD103+ DCs by gp96 represents an ideal vaccination strategy for priming effective and durable immunity in the epithelial tissues. It was previously shown that, based on differences in the localization and functions, there are 2 different subsets of lung TRM cells: airway TRM and interstitial TRM.^61-63^ CXCR6-CXCL16 interactions are crucial in controlling the localization of virus-specific TRM CD8+ T cells in the lungs and maintaining the airway TRM cell pool.^20^ Moreover, blocking of CXCR6-CXCL16 interactions significantly decreases the steady-state migration of TRM cells into airways, so vaccine induced SARS-CoV-2 S specific CD8+ T cells that express CXCR6 fulfill one of the major requirements for continued CXCR6 signaling in maintaining the airway TRM pool.^20^

We have confirmed in different infectious vaccine models that gp96-Ig carries all peptides of a cell that are selected in the recipient/vaccinee for MHC I, having the broadest, theoretically possible antigenic epitope-spectrum for cross-priming of CD8+ T cells by any MHC I type. Here, we showed that AD100-gp96-Ig-S resulted in the polyepitope and polyfunctional protein S-specific CD8+ and CD4+ T cell responses (Fig 3). When stimulated in vitro with S1+S2 peptides, spleen and lung CD8+ T cells produce IFNγ, TNFα and IL-2 cytokines (Fig 3) with CD8+ T cells in the lungs producing significantly less IFNγ than the CD8+ T cells in the spleen. It is known that enhanced activation, resulting from high levels of inflammation, induces CD8+ T cells entering the lungs to produce regulatory cytokines^64^ that initiate “dampening” of the immune response in order to prevent any excessive damage of the lung tissue. In addition, we report that CD4+ T cells in the lungs produce all 3 Th1 cytokines in equal ratio (Fig 3e). This finding is in line with our previous report discussing gp96-Ig induced SIV-specific CD4+ T cells in the lamina propria which were almost the exclusive producers of IL-2.^33^ Further studies will therefore be required to evaluate the role of protein-S specific CD4+ T cells in the induction of B cell and antibody responses. Previously, we have shown that gp96 is a powerful Th1 adjuvant for CTL priming and for stimulation of Th1 type antibodies that are of isotype IgG2a and IgG2b in mice and nonhuman primates.^37,38^ In addition, we will evaluate memory responses after a single and booster dose to establish the best vaccination protocol for future challenge studies.

In summary, we provide a paradigm for a novel vaccine development approach capable of induction of cellular immune responses in epithelial tissues such as the lungs. Structure-guided SARS-CoV-2 S protein combined with a safe and efficacious gp96-Ig vaccine platform can pave the way for a protective and durable immune response against COVID-19. This is a first demonstration of the utility and versatility of our proprietary secreted gp96-Ig SARS-CoV-2 vaccine platform that can be rapidly engineered and customized based on other and future pathogen sequences. Furthermore, the platform is proof of concept for the prototype vaccine approach for similar pathogens that require induction of effective TRM responses in epithelial tissues.

## Author Contributions

**NS** conceived and coordinated the experiments and obtained funding. **EF, LP, KP, KO**, and **NS** performed the experiments and analyzed the data. **MMS** provided reagents. **NS, EF, PJ, RJ**, and **MMS** wrote the paper. All authors were involved in writing and had final approval of the submitted and published versions of the paper.

## Acknowledgments

We dedicate this work to the late Dr. Eckhard Podack. We are grateful to all members of Strbo laboratory, Heat Biologics, Inc. CEO Jeff Wolf, Chief Scientific and Operating Officer, Jeff Hutchins, and Director Discovery Sciences, Eric Dixon, for their overall support, advice, and editorial contributions.

## References

1. Du L, He Y, Zhou Y, Liu S, Zheng BJ, Jiang S. The spike protein of SARS-CoV--a target for vaccine and therapeutic development. Nat Rev Microbiol. 2009;7(3):226–236.

2. Watanabe Y, Berndsen ZT, Raghwani J, et al. Vulnerabilities in coronavirus glycan shields despite extensive glycosylation. Nat Commun. 2020;11(1):2688.

3. Ibarrondo FJ, Fulcher JA, Goodman-Meza D, et al. Rapid decay of anti–SARS-CoV-2 antibodies in persons with mild Covid-19. N Engl J Med. 2020. doi: 10.1056/NEJMc2025179

4. Long QX, Tang XJ, Shi QL, et al. Clinical and immunological assessment of asymptomatic SARS-CoV-2 infections. Nat Med. 2020.

5. Channappanavar R, Fett C, Zhao J, Meyerholz DK, Perlman S. Virus-specific memory CD8 T cells provide substantial protection from lethal severe acute respiratory syndrome coronavirus infection. J Virol. 2014;88(19):11034–11044.

6. Tang F, Quan Y, Xin ZT, et al. Lack of peripheral memory B cell responses in recovered patients with severe acute respiratory syndrome: a six-year follow-up study. J Immunol. 2011;186(12):7264–7268.

7. Le Bert N, Tan AT, Kunasegaran K, et al. SARS-CoV-2-specific T cell immunity in cases of COVID-19 and SARS, and uninfected controls. Nature. 2020.

8. Andrew P Ferretti TK, Yifan Wang, Dalena MV Nguyen, Adam Weinheimer, Garrett S Dunlap, Qikai Xu, Nancy Nabilsi, Candace R Perullo, Alexander W Cristofaro, Holly J Whitton, Amy Virbasius, Kenneth J Olivier Jr., Lyndsey B Baiamonte, Angela T Alistar, Eric D Whitman, Sarah A Bertino, Shrikanta Chattopadhyay, Gavin MacBeath. COVID-19 Patients Form Memory CD8+ T Cells that Recognize a Small Set of Shared Immunodominant Epitopes in SARS-CoV-2. medRxiv. 2020.

9. Zhu N, Zhang D, Wang W, et al. A Novel Coronavirus from Patients with Pneumonia in China, 2019. New England Journal of Medicine. 2020;382(8):727–733.

10. Grifoni A, Weiskopf D, Ramirez SI, et al. Targets of T Cell Responses to SARS-CoV-2 Coronavirus in Humans with COVID-19 Disease and Unexposed Individuals. Cell. 2020;181(7):1489–1501 e1415.

11. Li CK, Wu H, Yan H, et al. T cell responses to whole SARS coronavirus in humans. J Immunol. 2008;181(8):5490–5500.

12. Zhao J, Zhao J, Perlman S. T cell responses are required for protection from clinical disease and for virus clearance in severe acute respiratory syndrome coronavirus-infected mice. J Virol. 2010;84(18):9318–9325.

13. Liu WJ, Lan J, Liu K, et al. Protective T Cell Responses Featured by Concordant Recognition of Middle East Respiratory Syndrome Coronavirus-Derived CD8+ T Cell Epitopes and Host MHC. J Immunol. 2017;198(2):873–882.

14. Peng Y, Mentzer AJ, Liu G, et al. Broad and strong memory CD4 (+) and CD8 (+) T cells induced by SARS-CoV-2 in UK convalescent COVID-19 patients. bioRxiv. 2020.

15. Altmann DM, Boyton RJ. SARS-CoV-2 T cell immunity: Specificity, function, durability, and role in protection. Sci Immunol. 2020;5(49).

16. Sekine T, Perez-Potti A, Rivera-Ballesteros O, et al. Robust T cell immunity in convalescent individuals with asymptomatic or mild COVID-19. bioRxiv. 2020.

17. Beura LK, Mitchell JS, Thompson EA, et al. Intravital mucosal imaging of CD8(+) resident memory T cells shows tissue-autonomous recall responses that amplify secondary memory. Nat Immunol. 2018;19(2):173–182.

18. Park SL, Zaid A, Hor JL, et al. Local proliferation maintains a stable pool of tissue-resident memory T cells after antiviral recall responses. Nat Immunol. 2018;19(2):183–191.

19. Wakim LM, Waithman J, van Rooijen N, Heath WR, Carbone FR. Dendritic cell-induced memory T cell activation in nonlymphoid tissues. Science. 2008;319(5860):198–202.

20. Wein AN, McMaster SR, Takamura S, et al. CXCR6 regulates localization of tissue-resident memory CD8 T cells to the airways. J Exp Med. 2019;216(12):2748–2762.

21. Agostini C, Cabrelle A, Calabrese F, et al. Role for CXCR6 and its ligand CXCL16 in the pathogenesis of T-cell alveolitis in sarcoidosis. Am J Respir Crit Care Med. 2005;172(10):1290–1298.

22. Freeman CM, Curtis JL, Chensue SW. CC chemokine receptor 5 and CXC chemokine receptor 6 expression by lung CD8+ cells correlates with chronic obstructive pulmonary disease severity. Am J Pathol. 2007;171(3):767–776.

23. Galkina E, Thatte J, Dabak V, Williams MB, Ley K, Braciale TJ. Preferential migration of effector CD8+ T cells into the interstitium of the normal lung. J Clin Invest. 2005;115(12):3473–3483.

24. Kohlmeier JE, Miller SC, Smith J, et al. The chemokine receptor CCR5 plays a key role in the early memory CD8+ T cell response to respiratory virus infections. Immunity. 2008;29(1):101–113.

25. Ray SJ, Franki SN, Pierce RH, et al. The collagen binding alpha1beta1 integrin VLA-1 regulates CD8 T cell-mediated immune protection against heterologous influenza infection. Immunity. 2004;20(2):167–179.

26. Slutter B, Pewe LL, Kaech SM, Harty JT. Lung airway-surveilling CXCR3(hi) memory CD8(+) T cells are critical for protection against influenza A virus. Immunity. 2013;39(5):939–948.

27. Hombrink P, Helbig C, Backer RA, et al. Programs for the persistence, vigilance and control of human CD8(+) lung-resident memory T cells. Nat Immunol. 2016;17(12):1467–1478.

28. Kumar BV, Ma W, Miron M, et al. Human Tissue-Resident Memory T Cells Are Defined by Core Transcriptional and Functional Signatures in Lymphoid and Mucosal Sites. Cell Rep. 2017;20(12):2921–2934.

29. Mackay LK, Rahimpour A, Ma JZ, et al. The developmental pathway for CD103(+)CD8+ tissue-resident memory T cells of skin. Nat Immunol. 2013;14(12):1294–1301.

30. Hogan RJ, Usherwood EJ, Zhong W, et al. Activated antigen-specific CD8+ T cells persist in the lungs following recovery from respiratory virus infections. J Immunol. 2001;166(3):1813–1822.

31. Wu T, Hu Y, Lee YT, et al. Lung-resident memory CD8 T cells (TRM) are indispensable for optimal cross-protection against pulmonary virus infection. J Leukoc Biol. 2014;95(2):215–224.

32. Zens KD, Chen JK, Farber DL. Vaccine-generated lung tissue-resident memory T cells provide heterosubtypic protection to influenza infection. JCI Insight. 2016;1(10).

33. Strbo N, Pahwa S, Kolber MA, Gonzalez L, Fisher E, Podack ER. Cell-secreted Gp96-Ig-peptide complexes induce lamina propria and intraepithelial CD8+ cytotoxic T lymphocytes in the intestinal mucosa. Mucosal Immunol. 2010;3(2):182–192.

34. Strbo N, Oizumi S, Sotosek-Tokmadzic V, Podack ER. Perforin is required for innate and adaptive immunity induced by heat shock protein gp96. Immunity. 2003;18(3):381–390.

35. Oizumi S, Strbo N, Pahwa S, Deyev V, Podack ER. Molecular and cellular requirements for enhanced antigen cross-presentation to CD8 cytotoxic T lymphocytes. J Immunol. 2007;179(4):2310–2317.

36. Selinger C, Strbo N, Gonzalez L, et al. Multiple low-dose challenges in a rhesus macaque AIDS vaccine trial result in an evolving host response that affects protective outcome. Clin Vaccine Immunol. 2014;21(12):1650–1660.

37. Strbo N, Garcia-Soto A, Schreiber TH, Podack ER. Secreted heat shock protein gp96-Ig: next-generation vaccines for cancer and infectious diseases. Immunol Res. 2013;57(1-3):311–325.

38. Strbo N, Vaccari M, Pahwa S, et al. Cutting edge: novel vaccination modality provides significant protection against mucosal infection by highly pathogenic simian immunodeficiency virus. J Immunol. 2013;190(6):2495–2499.

39. Strbo N, Vaccari M, Pahwa S, et al. Gp96 SIV Ig immunization induces potent polyepitope specific, multifunctional memory responses in rectal and vaginal mucosa. Vaccine. 2011;29(14):2619–2625.

40. Savaraj N, Wu CJ, Xu R, et al. Multidrug-resistant gene expression in small-cell lung cancer. Am J Clin Oncol. 1997;20(4):398–403.

41. Yamazaki K, Spruill G, Rhoderick J, Spielman J, Savaraj N, Podack ER. Small cell lung carcinomas express shared and private tumor antigens presented by HLA-A1 or HLA-A2. Cancer Res. 1999;59(18):4642–4650.

42. Morgensztern D WS, Bazhenova L, McDermott L, Hutchins J, Yalor DH, Robinson FL, Dowdell AK, Piening BD, Harb W, Pannell N, Cohen RB. Tumor antigen expression and survival of patients with previously-treated advanced NSCLC recieving viagenpumatucel-L (HS-110) plus nivolumab. Presented at: ASCO;

43. Yamazaki K, Nguyen T, Podack ER. Cutting edge: tumor secreted heat shock-fusion protein elicits CD8 cells for rejection. J Immunol. 1999;163(10):5178–5182.

44. Schenkel JM, Masopust D. Tissue-resident memory T cells. Immunity. 2014;41(6):886–897.

45. Masopust D, Vezys V, Usherwood EJ, et al. Activated primary and memory CD8 T cells migrate to nonlymphoid tissues regardless of site of activation or tissue of origin. J Immunol. 2004;172(8):4875–4882.

46. Gonzalez L, Strbo N, Podack ER. Humanized mice: novel model for studying mechanisms of human immune-based therapies. Immunol Res. 2013;57(1-3):326–334.

47. Vaccari M, Gordon SN, Fourati S, et al. Adjuvant-dependent innate and adaptive immune signatures of risk of SIVmac251 acquisition. Nat Med. 2016;22(7):762–770.

48. van den Brand JM, Haagmans BL, van Riel D, Osterhaus AD, Kuiken T. The pathology and pathogenesis of experimental severe acute respiratory syndrome and influenza in animal models. J Comp Pathol. 2014;151(1):83–112.

49. Kropp LE, Garg M, Binder RJ. Ovalbumin-derived precursor peptides are transferred sequentially from gp96 and calreticulin to MHC class I in the endoplasmic reticulum. J Immunol. 2010;184(10):5619–5627.

50. Srivastava P. Roles of heat-shock proteins in innate and adaptive immunity. Nat Rev Immunol. 2002;2(3):185–194.

51. Binder RJ, Srivastava PK. Essential role of CD91 in re-presentation of gp96-chaperoned peptides. Proc Natl Acad Sci U S A. 2004;101(16):6128–6133.

52. Messmer MN, Pasmowitz J, Kropp LE, Watkins SC, Binder RJ. Identification of the cellular sentinels for native immunogenic heat shock proteins in vivo. J Immunol. 2013;191(8):4456–4465.

53. Yang ZY, Kong WP, Huang Y, et al. A DNA vaccine induces SARS coronavirus neutralization and protective immunity in mice. Nature. 2004;428(6982):561–564.

54. Lai MMC, and K. V. Holmes. Fields Virology. Philadelphia, PA.: Lippincott Williams & Wilkins; 2001.

55. Ni L, Ye F, Cheng ML, et al. Detection of SARS-CoV-2-Specific Humoral and Cellular Immunity in COVID-19 Convalescent Individuals. Immunity. 2020;52(6):971–977 e973.

56. Shane HL, Klonowski KD. Every breath you take: the impact of environment on resident memory CD8 T cells in the lung. Front Immunol. 2014;5:320.

57. Liao M, Liu Y, Yuan J, et al. Single-cell landscape of bronchoalveolar immune cells in patients with COVID-19. Nat Med. 2020;26(6):842–844.

58. Belyakov IM, Hammond SA, Ahlers JD, Glenn GM, Berzofsky JA. Transcutaneous immunization induces mucosal CTLs and protective immunity by migration of primed skin dendritic cells. J Clin Invest. 2004;113(7):998–1007.

59. del Rio ML, Bernhardt G, Rodriguez-Barbosa JI, Forster R. Development and functional specialization of CD103+ dendritic cells. Immunol Rev. 2010;234(1):268–281.

60. Bedoui S, Whitney PG, Waithman J, et al. Cross-presentation of viral and self antigens by skin-derived CD103+ dendritic cells. Nat Immunol. 2009;10(5):488–495.

61. Jozwik A, Habibi MS, Paras A, et al. RSV-specific airway resident memory CD8+ T cells and differential disease severity after experimental human infection. Nat Commun. 2015;6:10224.

62. McMaster SR, Wilson JJ, Wang H, Kohlmeier JE. Airway-Resident Memory CD8 T Cells Provide Antigen-Specific Protection against Respiratory Virus Challenge through Rapid IFN-gamma Production. J Immunol. 2015;195(1):203–209.

63. Zhao J, Zhao J, Mangalam AK, et al. Airway Memory CD4(+) T Cells Mediate Protective Immunity against Emerging Respiratory Coronaviruses. Immunity. 2016;44(6):1379–1391.

64. Sun J, Dodd H, Moser EK, Sharma R, Braciale TJ. CD4+ T cell help and innate-derived IL-27 induce Blimp-1-dependent IL-10 production by antiviral CTLs. Nat Immunol. 2011;12(4):327–334.

